# A symbiont-derived biosurfactant couples bacterial surface properties to host-mediated clearance

**DOI:** 10.64898/2026.09.18.752026

**Authors:** Barbara Pees, Celia Escudero-Hernández, Avril von Hoyningen-Huene, Philip Rosenstiel, Ruth A. Schmitz, Katja Dierking

**Affiliations:** Department of Evolutionary Ecology and Genetics, Zoological Institute, Christian Albrechts University, Kiel, Germany; Department of Paediatrics and Immunology, University of Valladolid, Valladolid, Spain; Institute of Clinical Molecular Biology, Christian Albrechts University and University Hospital Schleswig-Holstein, Kiel, Germany; Institute for General Microbiology, Christian Albrechts University, Kiel, Germany; Helmholtz Institute for Functional Marine Biodiversity at the University of Oldenburg (HIFMB), Oldenburg, Germany; Alfred Wegener Institute Helmholtz Centre for Polar and Marine Research, Bremerhaven, Germany

## Abstract

Host mechanisms that regulate intestinal colonization are viewed as acting on microbes, yet microbial traits that determine susceptibility to host control remain poorly understood. Using the association between *Caenorhabditis elegans* and its symbiont *Pseudomonas lurida* MYb11, we identify the cyclic lipopeptide biosurfactant massetolide as a bacterial factor that couples microbial physiology to host-mediated clearance. An unbiased screen of >11,000 bacterial mutants coupled to a host transcriptional reporter, combined with bacterial and host genetics, chemical complementation, transcriptomics, and physiological analyses, revealed that massetolide triggers host TGF-β and serotonergic signaling to promote intestinal peristalsis, limiting colonization. Massetolide also alters bacterial surface properties and promotes swarming. Experimentally increased serotonin-dependent peristalsis selectively reduces colonization by massetolide-producing bacteria, whereas massetolide-deficient bacteria remain resistant to host-mediated expulsion. Together, our findings reveal a symbiont-derived biosurfactant that links host neuroimmune control of intestinal peristalsis with bacterial susceptibility to expulsion, coupling host and microbial physiology to regulate symbiont abundance.

## Introduction

A central challenge of host–microbe symbiosis is how hosts maintain beneficial symbionts while preventing their uncontrolled expansion that is often linked to host exploitation and subsequent host fitness reductions. Colonization depends not only on host responses but also on microbial traits that influence susceptibility to host control^1,2^. Although considerable progress has been made in identifying the molecular mechanisms underlying host colonization^3–5^, the specific bacterial stimuli detected by the host and the precise coordination of host responses required to control bacterial colonization remain poorly understood.

One potential bacteria-derived cue are biosurfactants, widespread microbial secondary metabolites that regulate bacterial surface behavior, competition, and community dynamics^6,7^. Beyond these microbial functions, accumulating evidence suggests that biosurfactants can also influence host physiology. For example, cyclic lipopeptides from pathogenic *Bacillus* spp. and *Pseudomonas* spp. can induce systemic resistance in plants^8^, indicating that biosurfactants can function as host-directed signals in the defense against pathogens. However, whether biosurfactants modulate host physiology in animals and, more specifically, the association between animals and their non-pathogenic symbionts remain largely unexplored.

Here, we use the facultative association between *C. elegans* and its natural symbiont *Pseudomonas lurida* MYb11 to study both bacterial signals and host responses that shape colonization. Combining an unbiased bacterial genetic screen with a host-response reporter and performing bacterial and host functional analyses, we identified the cyclic lipopeptide biosurfactant massetolide as a key mediator of colonization through a dual mechanism: First, massetolide is used as a signal by the host, initiating TGF-β and serotonergic signaling pathways to promote intestinal peristalsis and bacterial expulsion. Second, massetolide enables bacteria to swarm, but also renders them susceptible to serotonin-dependent clearance in the context of host association. Our findings uncover that a bacterial biosurfactant couples host and microbial physiology to regulate colonization.

## Results

### *P. lurida* MYb11 strongly induces expression of the *C. elegans* gene *mbug-1*

*P. lurida* MYb11 is a natural *C. elegans* symbiont which protects the nematode against infection with the bacterial pathogens *Bacillus thuringiensis* and *Pseudomonas aeruginosa* and with the intracellular microsporidian pathogen *Nematocida parisii*. MYb11 produces two cyclic lipopeptide biosurfactants of the viscosin group, massetolide E and massetolide F, that directly inhibit pathogen growth^9,10^. Additionally, MYb11 activates host defense responses^11^, though the identity of the symbiont-derived molecule that activates these host responses and the function of this host activation was unknown.

Part of the *C. elegans* response triggered by MYb11 involves the upregulation of the gene F55G11.4^11^. The gene encodes a nematode specific CUB-domain containing protein of unknown function, which has previously been reported as pathogen-responsive gene^12–15^. We monitored gene expression *in vivo* using a F55G11.4p::GFP transcriptional reporter^11^ and observed that the MYb11-induced expression of F55G11.4 exceeds induction by the *C. elegans* pathogen *P. aeruginosa* PA14 by 8-fold (Figure 1a, c). The Gram-positive pathogens *B. thuringiensis* Bt679 and Bt247 did not induce expression (Figure 1b; Supplemental Figure 1). Thus, we named F55G11.4 <u>m</u>icro<u>b</u>iota-<u>u</u>pregulated <u>g</u>ene 1, *mbug-1*. As previously reported^11^, MYb11 most strongly induces *mbug-1* expression in the first intestinal ring (Figure 1d). Here, we additionally observed that MYb11-induced *mbug-1* expression extends to the whole intestine and the epidermis in older adults (Figure 1e, f).

**Figure 1.**
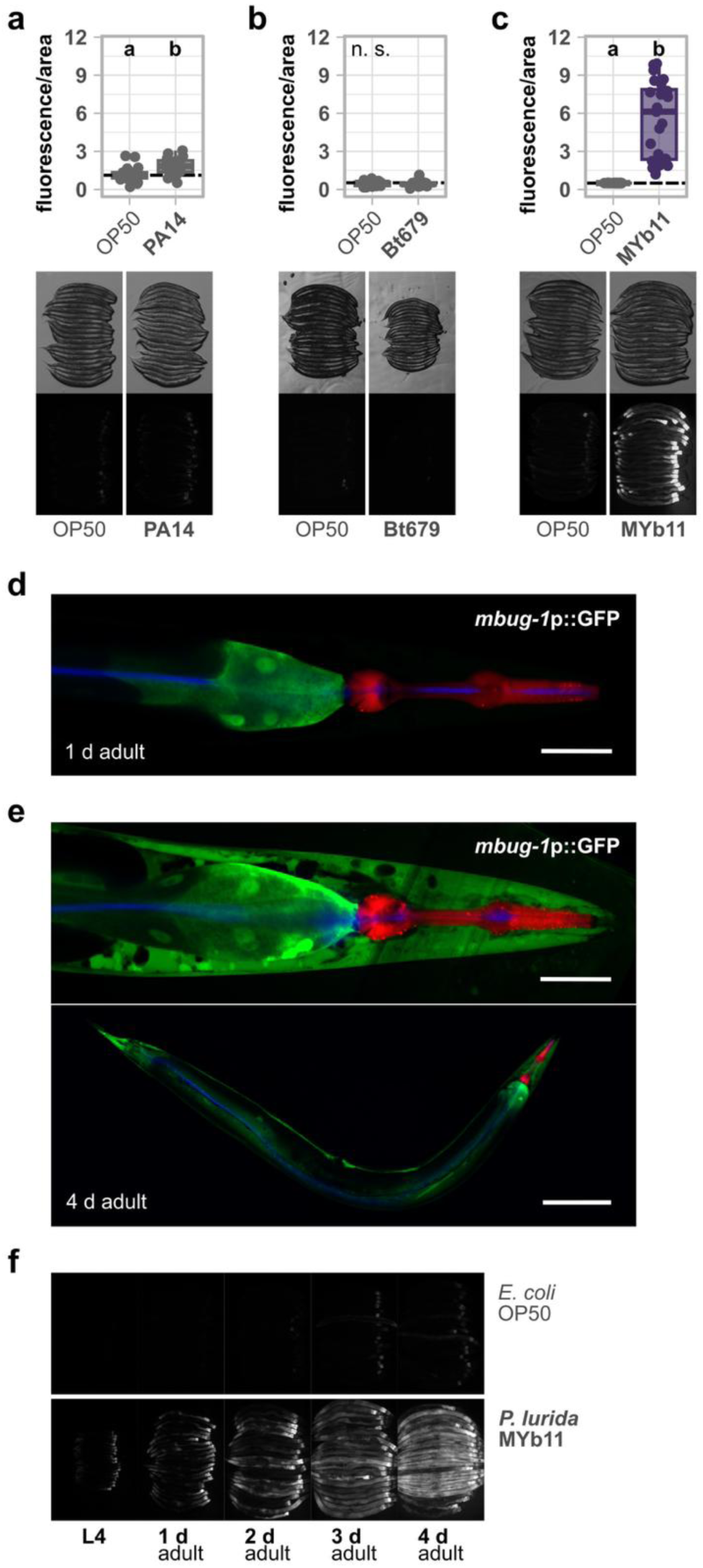
*P. lurida* MYb11 strongly induces *mbug-1* expression in the intestine and the epidermis. (a-c) Quantification of *mbug-1*p::GFP fluorescence in 1 d adult transgenic worms exposed to *E. coli* OP50 and pathogenic (a) *P. aeruginosa* PA14, (b) *B. thuringiensis* Bt679 or symbiont (c) MYb11. Fluorescence was normalized by the worm’s body size (area). Each dot represents one worm with n = 20–25, and the dashed line represents the median of the mean gray value for OP50-exposed worms. Statistical significance (*p* ≤ 0.05) among groups of differently exposed worms was assessed using the Wilcoxon rank sum test. Different letters indicate significant differences, whereas shared letters indicate no significant difference. Representative images of groups of 20 individuals, arranged with their heads pointing to the right, are shown below in bright field (top) and fluorescence (bottom). (d, e) Microscopic images of a transgenic strain carrying *mbug-1*p::GFP (green), *myo-2*p::mCherry (red) exposed to MYb11::CFP (blue). Shown are the head region including the first intestinal cells of a 1 d and 4 d adult as well as the full body of a 4 d adult. Scale bars represent 50 µm and (e) 200 µm (bottom). (f) Expression of *mbug-1*p::GFP over time in worms exposed to OP50 or MYb11. Fluorescence images of groups of 20 individuals, arranged with their heads pointing to the right, are shown. Raw data and corresponding *p*-values are provided in Extended Data Table 1 and Supplementary Table 1, respectively.

### A comprehensive *P. lurida* MYb11 mutant screen identified massetolide to upregulate *mbug-1*

In order to investigate symbiont-derived molecules that trigger the *C. elegans* response, we first tested whether the bacterial factor that induces *mbug-1* expression is secreted. We mixed *E. coli* OP50 with filter-sterilized supernatant of a MYb11 overnight culture and monitored *mbug-1* expression in the transcriptional reporter. The MYb11 supernatant was sufficient to induce *mbug-1* expression *in vivo*, although less strongly than continuous exposure to MYb11 (Figure 2a), suggesting that a secreted MYb11 compound mediates the host response.

**Figure 2.**
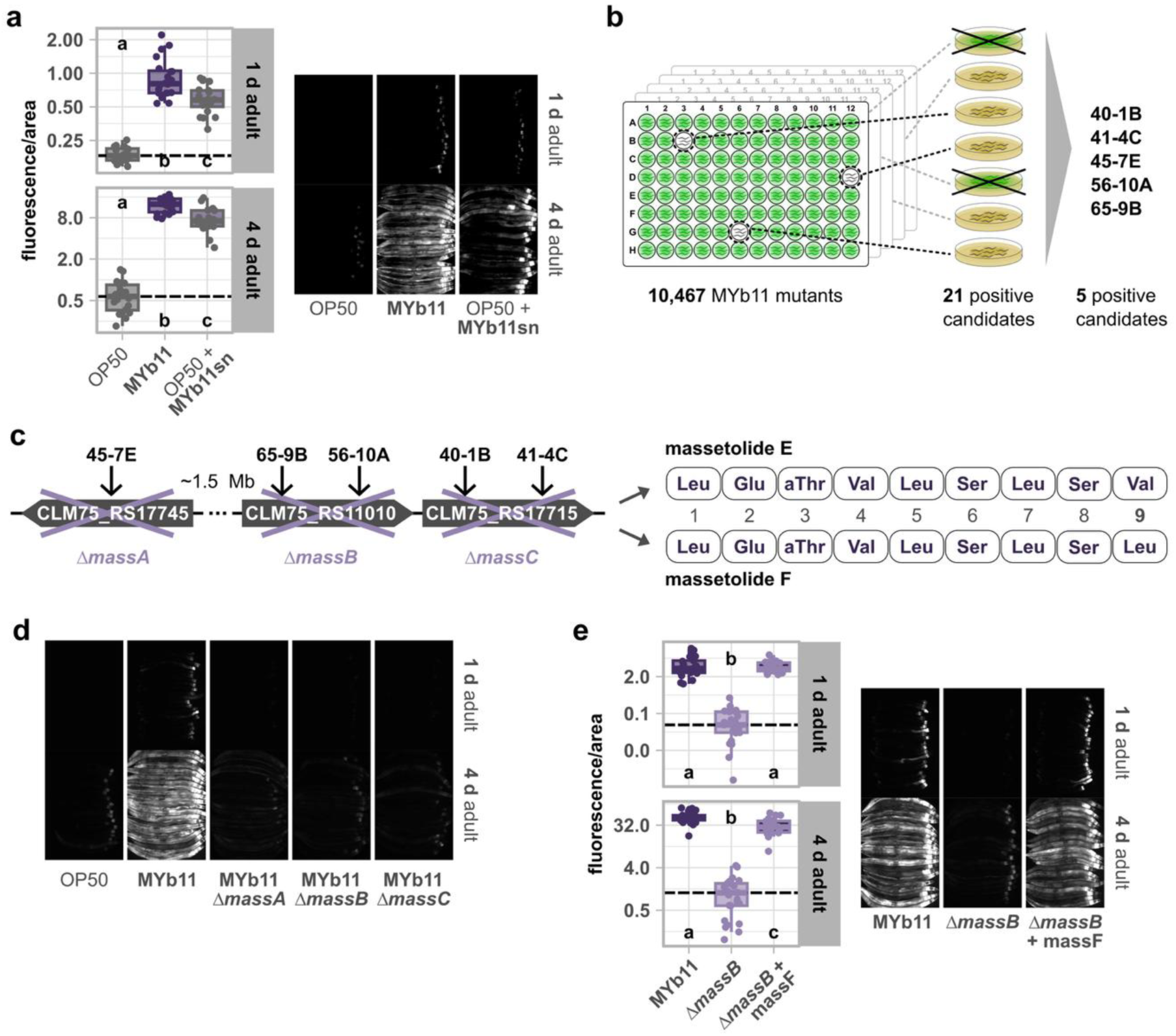
MYb11-derived massetolide induces *mbug-1* expression in the intestine and the epidermis. (a, e) Quantification of *mbug-1*p::GFP fluorescence in 1 d and 4 d adult transgenic worms exposed to (a) *E. coli* OP50, MYb11, or OP50 cells mixed with sterile, cell-free MYb11 supernatant (MYb11sn), or (e) MYb11, MYb11 *ΔmassB*, or MYb11 *ΔmassB* supplemented with 2 µg/µL purified massetolide F in the inoculum. Fluorescence was normalized by the worm’s body size (area). Each dot represents one worm with (a) n = 20 or (e) n = 25, and the dashed line represents the median of the mean gray value for (a) OP50-exposed or (e) *ΔmassB*-exposed worms. Statistical significance (*p* ≤ 0.05) among groups of differently exposed worms was assessed using the Kruskal–Wallis rank sum test followed by Dunn’s *post hoc* test with Holm correction. Different letters indicate significant differences, whereas shared letters indicate no significant difference. Fluorescence images of groups of 20 individuals, arranged with their heads pointing to the right, are shown on the right. (b) Schematic of MYb11 transposon mutant library screen. (c) Biosynthetic gene cluster producing massetolide E and F. The approximate transposon sites of the candidate strains derived from the transposon mutant screen are shown as arrows. (d) Expression of *mbug-1*p::GFP in 1 d and 4 d adults exposed to OP50, MYb11, or three massetolide-deficient MYb11 mutants, *ΔmassA*, *ΔmassB*, and *ΔmassC*. Representative fluorescence images of groups of 20 individuals, arranged with their heads pointing to the right, are shown. Raw data and corresponding *p*-values are provided in Extended Data Table 1 and Supplementary Table 1, respectively.

To identify the MYb11-derived, secreted compound that triggers the host response, we generated and screened a MYb11 transposon mutant library consisting of approx. 11,000 transposon mutants for mutants unable to induce *mbug-1* expression (Figure 2b). Following a pre-screen that yielded 21 potential candidates, we identified five transposon mutants which failed to induce *mbug-1* expression (Supplemental Figure 2a). We detected the transposon insertions to be located in the three genes of the massetolide biosynthetic gene cluster, encoding the ribosomal peptide synthetases (NRPS) genes CLM75_RS17745, CLM75_RS11010, and CLM75_RS11015 that are located at two different loci in the MYb11 genome. The MYb11 massetolide NRPS genes, designated *massA*, *massB*, and *massC*, produce massetolide E and F (Figure 2c)^9,16,17^, which are structural analogs that differ by a single amino acid residue at position 9 of their peptide sequence^16^. To validate our reporter screen, we generated MYb11 mutants with individual, paired, or complete knockouts of the three NRPS genes *massA*, *massB*, and *massC*, and exposed the *mbug-1* reporter to the NRPS-deficient mutants. We observed that already the MYb11 *ΔmassA, ΔmassB, and ΔmassC* single mutants were not able to induce *mbug-1* expression (Figure 2d; Supplemental Figure 2b-d).

Finally, we employed purified massetolide F that was previously identified by comparative mass spectrometry using MYb11 and MYb11 *Δmass* mutants, and then isolated and characterized using preparative HPLC and NMR spectroscopy, respectively (co-submitted study Wang et al.). Indeed, adding purified massetolide F to *ΔmassB* mutants rescued the expression of *mbug-1* in 1 d and 4 d adults (Figure 2e; Supplemental Figure 2e).

These observations demonstrate that MYb11-derived massetolide, which was previously identified to act as antimicrobial compound in host protection against pathogens^9,10^, also acts as host-directed signal and induces a gene expression response in *C. elegans*.

### Massetolide modulates bacterial physiology and host phenotypes

Bacterial biosurfactants are increasingly recognized as multifunctional ecological mediators that are crucial for microbial physiology and ecology, particularly influencing environmental colonization and microbial competition^7^. However, biosurfactant-mediated, direct communication with the host is much less systematically studied. Hence, we used our massetolide-deficient mutants to explore the function of massetolide in MYb11 physiology and in the interaction with the *C. elegans* host. First, we tested the predication that massetolide acts as a biosurfactant. The MYb11 *Δmass* mutants kept their dome-shaped drop on a hydrophobic surface and failed to swarm on a low-agar surface, unlike wildtype MYb11, demonstrating the surface-active properties of massetolide (Figure 3a).

**Figure 3.**
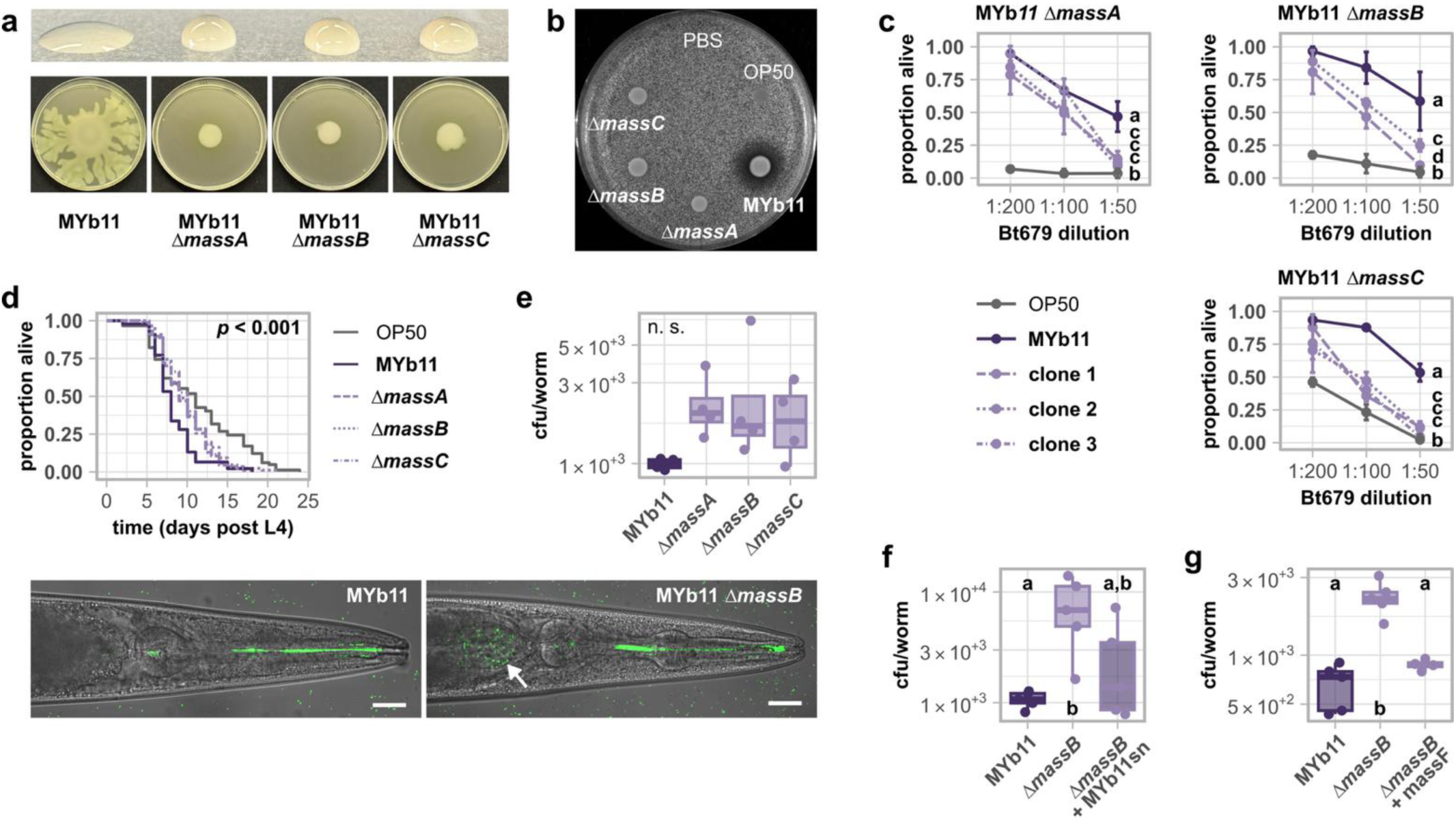
Massetolide modulates bacterial physiology and host phenotypes. (a) Photographs showing drops of overnight cultures of MYb11, and the massetolide-deficient MYb11 *Δmass* mutants on a hydrophobic surface (top) and swarming ability of the same bacteria on low concentration agar plates (bottom). (b) Photograph showing the inhibition zone formed by MYb11 or the *Δmass* mutants against *B. thuringiensis* Bt679 incorporated into the agar. Phosphate-buffered saline (PBS) served as buffer control. (c) Survival of wildtype N2 infected with serial dilutions of *B. thuringiensis* Bt679 after 24 hours post infection (hpi). Worms were fed with either *E. coli* OP50, MYb11, or clones of massetolide mutants before and during infection. Each dot represents the mean ± standard deviation of three worm populations, n = 3. The same letters indicate no significant differences between the dose-response curves according to a generalized linear model followed by Tukey-adjusted *post hoc* multiple comparisons. (d) Lifespan of wildtype N2 exposed to OP50, MYb11, or massetolide-deficient MYb11 mutants shown as Kaplan-Meier curves. Statistical significance (p ≤ 0.05) between treatment groups was determined using a Cox proportional hazards model with replicate populations of 30 individuals as frailty term, n = 3. (e-g) Bacterial load of MYb11, massetolide-deficient MYb11 *Δmass* mutants, (f) MYb11 *ΔmassB* cells mixed with sterile, cell-free MYb11 supernatant (MYb11sn), or (g) MYb11 *ΔmassB* supplemented with 2 µg/µL purified massetolide F in 1 d adults. Statistical significance (*p* ≤ 0.05) among groups of differently exposed worms was assessed using the Kruskal–Wallis rank sum test followed by Dunn’s *post hoc* test with Holm correction with n = 4-5. (e) Microscopic images display 1 d adults fed with MYb11::GFP or MYb11 *ΔmassB*::GFP (green). Shown is the head region including the first intestinal cells. The arrow points to an accumulation of MYb11 *ΔmassB* in the first intestinal ring. Scale bars represent 20 µm. Raw data and corresponding *p*-values are provided in Extended Data Table 1 and Supplementary Table 1, respectively.

Next, we confirmed the antimicrobial activity of massetolide and tested MYb11 *Δmass* mutants for their ability to inhibit growth of the Gram-positive *C. elegans* pathogen *B. thuringiensis* Bt679 *in vitro*. We found that all mutants had lost the antimicrobial activity against Bt679 (Figure 3b). These results validate our previous work, demonstrating that massetolide has antimicrobial properties, inhibiting growth of Gram-positive bacteria, including *B. thuringiensis* and *Staphylococcus aureus*^9^.

We then tested if massetolide production is also required for the protection of *C. elegans* against *B. thuringiensis* Bt679 infection *in vivo*, as previously assumed^9^. Interestingly, MYb11-mediated protection against Bt679 infection was only partially reduced (Figure 3c), even in worms exposed to MYb11 *Δmass* double/triple mutants (Supplemental Figure 3a-d), suggesting the presence of additional, yet unknown protective mechanisms that may depend on microbe-host interactions. Besides its beneficial function in protecting the host against pathogen infection^9,18^, MYb11 negatively affects *C. elegans* lifespan^11,19,20^. We now found that the absence of massetolide partially rescued the lifespan decrease of worms exposed to MYb11 wildtype (Figure 3d). This result indicates that massetolide may have toxic effects during *C. elegans* ageing, a process that is associated with increased intestinal bacterial proliferation^21,22^. This observation also underscores the ambivalent role of MYb11 in the interaction with the host – protective during infection, but decreasing lifespan under non-infectious conditions.

Finally, we wondered whether MYb11-derived massetolide affects bacterial colonization of the *C. elegans* gut. To this end, we isolated bacteria from the *C. elegans* gut and counted colony forming units (cfu) and, additionally, observed colonization using fluorescent strains of MYb11 wildtype and the massetolide-deficient MYb11 *ΔmassB* mutant. We found that MYb11 *Δmass* mutants colonized the worm gut better than MYb11 wildtype bacteria (Figure 3e) despite their similar growth dynamics *in vitro* (Supplemental Figure 3e). Furthermore, not only filter-sterilized MYb11 supernatant containing massetolide is sufficient to rescue the high colonization of *C. elegans* by the MYb11 *ΔmassB* mutant (Figure 3f), but also supplementation of purified massetolide F (Figure 3g).

In summary, we discovered that MYb11-derived massetolide has far-reaching effects on both, bacterial physiology and the host: massetolide has surface-active properties and is required for bacterial mobility and antimicrobial activity. On the host side, MYb11-mediated protection against *B. thuringiensis* Bt679 infection partially depends on massetolide production. Furthermore, MYb11-derived massetolide reduces *C. elegans* lifespan and bacterial colonization of the *C. elegans* gut.

### Massetolide affects *P. lurida* MYb11 colonization through DBL-1/TGF-β signaling

We found that the massetolide-deficient MYb11 *Δmass* mutants exhibit enhanced colonization while the supplementation of purified massetolide reduced colonization to MYb11 wildtype level. This is an intriguing finding, since it indicates that massetolide may act as a symbiont-derived signal that limits bacterial colonization within the host, likely through activating *C. elegans* regulatory pathways.

Transforming Growth Factor-beta/Bone Morphogenetic Protein (TGF-β/BMP) immune signaling has been implicated in regulating bacterial colonization of the *C. elegans* intestine by its natural microbiota^23,24^. To investigate a potential link between MYb11-derived massetolide and TGF-β signaling, we assessed MYb11 colonization in *C. elegans* mutants of the TGF-β Sma/Mab pathway, targeting the TGF-β/BMP ligand encoding gene *dbl-1*, type I receptor encoding gene *sma-6*, the R-Smad encoding genes *sma-2* and *sma-3*, and Co-Smad encoding gene *sma-4* (Figure 4a). MYb11 colonization was significantly increased in *C. elegans* mutants *dbl-1(nk3)*, *sma-6(e1482)*, *sma-4(e729)* (all ∼100-fold), *sma-2(e502)* (∼50-fold), and *sma-3(e491)* (∼10-fold) compared to wildtype *C. elegans* N2 (Figure 4b-d).

**Figure 4.**
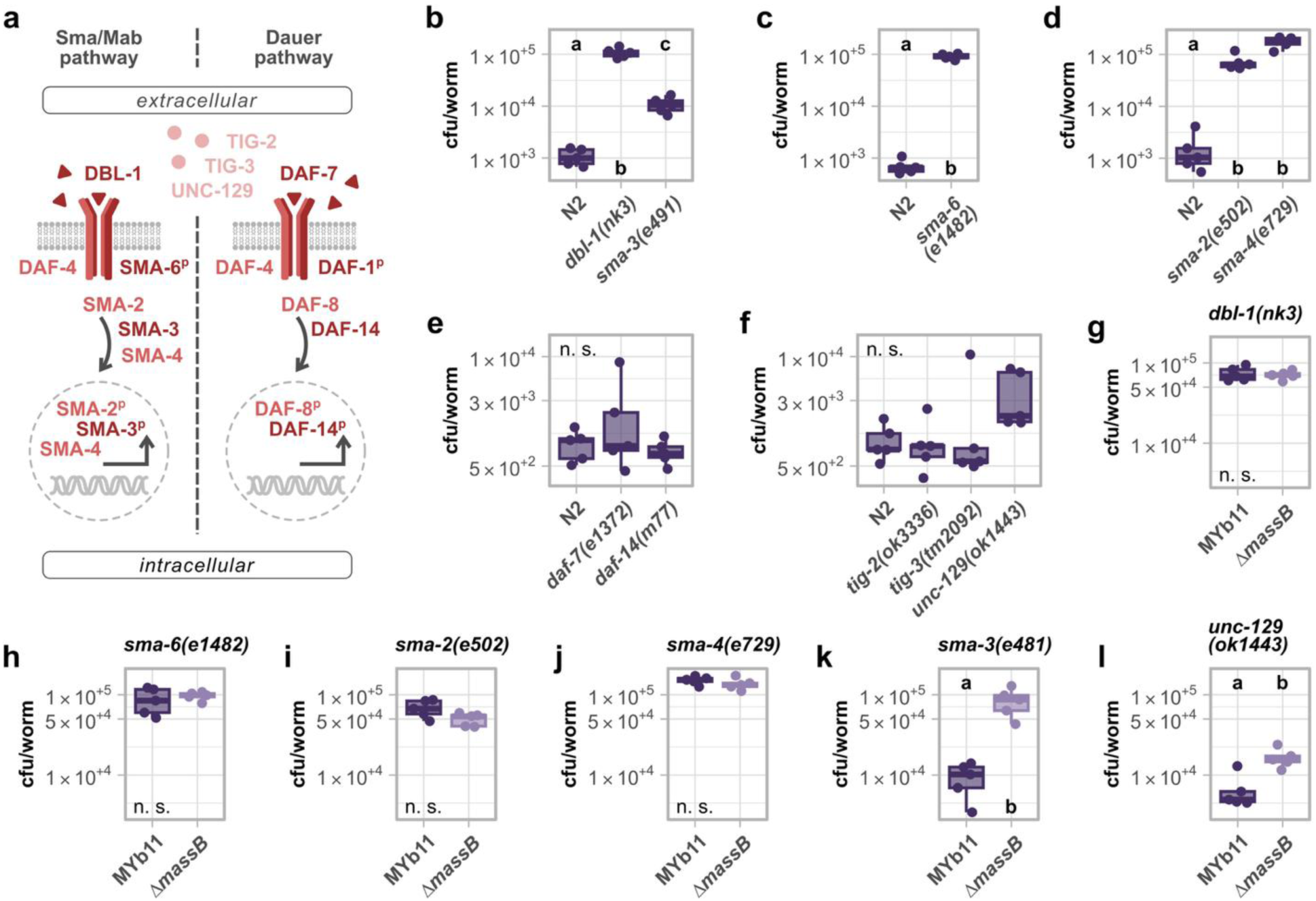
Massetolide affects *P. lurida* MYb11 colonization through DBL-1/TGF*-*β signaling. (a) Schematic of TGF-β signaling showing the Sma/Mab, the Dauer pathway, and orphan ligands TIG-2, TIG-3, and UNC-129. Adapted from^28^. (b-l) Bacterial load of (b-f) MYb11, or (g-l) MYb11 and the massetolide-deficient MYb11 *ΔmassB* mutant in wildtype N2 and TGF-β signaling mutants. Statistical significance (*p* ≤ 0.05) among treatment groups was assessed either using (b, d-f) the Kruskal–Wallis rank sum test followed by Dunn’s *post hoc* test with Holm correction or using (c, g-l) the Wilcoxon rank sum test with n = 5. Raw data and corresponding *p*-values are provided in Extended Data Table 1 and Supplementary Table 1, respectively.

In addition to *dbl-1*, the *C. elegans* genome contains four genes encoding TGF-β-related ligands: *daf-7*, *tig-2*, *tig-3*, and *unc-129* (Figure 4a). DAF-7 is more related to the TGF-β/Activin branch of the TGF-β superfamily and plays an important role in dauer diapause, an alternative *C. elegans* developmental program, lifespan, and lipid storage^25^. The co-submitted study by Wang *et al.* demonstrates that MYb11-derived massetolide affects *C. elegans* dauer formation through insulin signaling and the DAF-7/TGF-β pathway. We thus tested mutants of the DAF-7/TGF-β pathway, targeting TGF-β ligand encoding gene *daf-7* and type II receptor encoding gene *daf-14* in the context of host colonization, but neither mutant showed increased colonization by MYb11 (Figure 4e). We also tested the involvement of the three remaining TGF-β orphan ligands, TIG-2, TIG-3, and UNC-129 that play a role in directing axon growth^26,27^ for their involvement in intestinal colonization by MYb11. While we did not find any deviating colonization phenotype in *tig-2(ok3336)* and *tig-3(tm2092)* mutants, colonization was slightly increased in the *unc-129(ok1443)* mutant (Figure 4f).

We next wanted to understand whether the effect of MYb11-derived massetolide on host colonization (reduction in colonization) is mediated by TGF-β signaling. Hence, we assessed colonization of *C. elegans dbl-1*, *sma-6*, *sma-2*, *sma-3*, *sma-4*, and *unc-129* mutants by MYb11 wildtype and *ΔmassB* mutants. In *C. elegans dbl-1(nk3)*, *sma-6(e1482)*, *sma-2(e502)*, and *sma-4(e729)* mutants, there was no difference between colonization by massetolide-producing MYb11 wildtype and massetolide-deficient MYb11 *ΔmassB* mutants (Figure 4g-j; Supplemental Figure 4a, b), indicating that massetolide acts through DBL-1/TGF-β signaling, including the canonical receptor SMA-6, to limit bacterial colonization by MYb11. In contrast, in *C. elegans sma-3(e491)* and *unc-129(ok1443)* mutants, the difference between colonization by MYb11 wildtype and MYb11 *ΔmassB* mutants remained (Figure 4k, l; Supplemental Figure 4c), indicating that *unc-129* and *sma-3* regulate host colonization independently of massetolide.

These findings suggest that a non-canonical DBL-1/TGF-β signaling pathway mediates control of host colonization through massetolide.

### Serotonin acts downstream of DBL-1/TGF-β signaling to regulate *P. lurida* MYb11 colonization

Our results suggest that massetolide signals through a non-canonical DBL-1/TGF-β signaling pathway to control colonization. To clarify the downstream mechanism, we firstly reanalyzed a published RNA-seq dataset including the *C. elegans dbl-1(nk3)* mutant and a *dbl-1* overexpression strain^29^ to identify genes that are positively regulated by TGF-β signaling, for example genes downregulated in the knockout mutants and upregulated in the overexpression strain, relative to wildtype N2 (Figure 5a). Additionally, we generated our own RNA-seq data comparing the *C. elegans* response to the massetolide-producing MYb11 wildtype against the massetolide-deficient *Δmass* mutants to identify host genes responsive to massetolide. We then examined the expression of the identified 47 DBL-1/TGF-β target genes in our own RNA-seq data. We found one gene, whose expression was also dependent on massetolide (i. e. activation of expression by massetolide-producing wildtype MYb11, but not by massetolide-deficient *Δmass* mutants or the food bacterium *E. coli* OP50): *bas-1* (Figure 5b). We confirmed that massetolide induces *bas-1* expression *in vivo* using the *C. elegans* reporter construct BAS-1::GFP, which showed higher levels of GFP-tagged BAS-1 in worms exposed to MYb11 wildtype compared to those exposed to massetolide-deficient mutant *ΔmassB* or OP50 (Figure 5c). Interestingly, the increased abundance of BAS-1::GFP in worms exposed to MYb11 wildtype was primarily localized to the first intestinal cells (Figure 5c, d). Next, we tested if *bas-1* is required for the regulation of *C. elegans* colonization by MYb11 and assessed bacterial load in the *C. elegans bas-1(ad446)* and *bas-1(tm351)* mutants. We found that colonization by MYb11 was significantly increased in both *bas-1* mutants (Figure 5e; Supplemental Figure 5a). Importantly, colonization of the *bas-1* mutants was unaffected by massetolide production in MYb11 strains (Figure 5f). This indicates that BAS-1 function is required for the massetolide-dependent colonization phenotype. The gene *bas-1* encodes an aromatic amino acid decarboxylase required for converting levadopa (L-DOPA) to dopamine and 5-hydroxytryptophan (5-HTP) to 5-hydroxytryptamine (5-HT, serotonin)^30^. Hence, we hypothesized that either dopamine or serotonin influences massetolide-dependent control of *C. elegans* colonization by MYb11 downstream of TGF-β signaling. To test this hypothesis, we utilized *C. elegans* mutants of the rate-limiting enzymes for dopamine, the tyrosine hydroxylase encoding gene *cat-2*, and serotonin, namely the tryptophan hydroxylase encoding gene *tph-1*, and assessed MYb11 colonization. We found that MYb11 colonization was significantly increased in both mutants, *cat-2(n4547)* and *tph-1(mg280)* (Figure 5g; Supplemental Figure 5b). Also, massetolide production did not influence the colonization of *cat-2* and *tph-1*, as massetolide-producing MYb11 wildtype and massetolide-deficient MYb11 *ΔmassB* mutants showed comparable colonization levels (Figure 5h, i). Exogenous serotonin restored the high colonization phenotype of the *tph-1* mutant, but intriguingly, only for colonization by MYb11 wildtype, not by the *ΔmassB* mutant (Figure 5j, k). In contrast, exogenous dopamine, if at all, had only a very slight effect in reducing the high colonization of the *cat-2* mutant (Supplemental Figure 5c). These results suggest that serotonin in particular may play a role in regulating MYb11 colonization.

**Figure 5.**
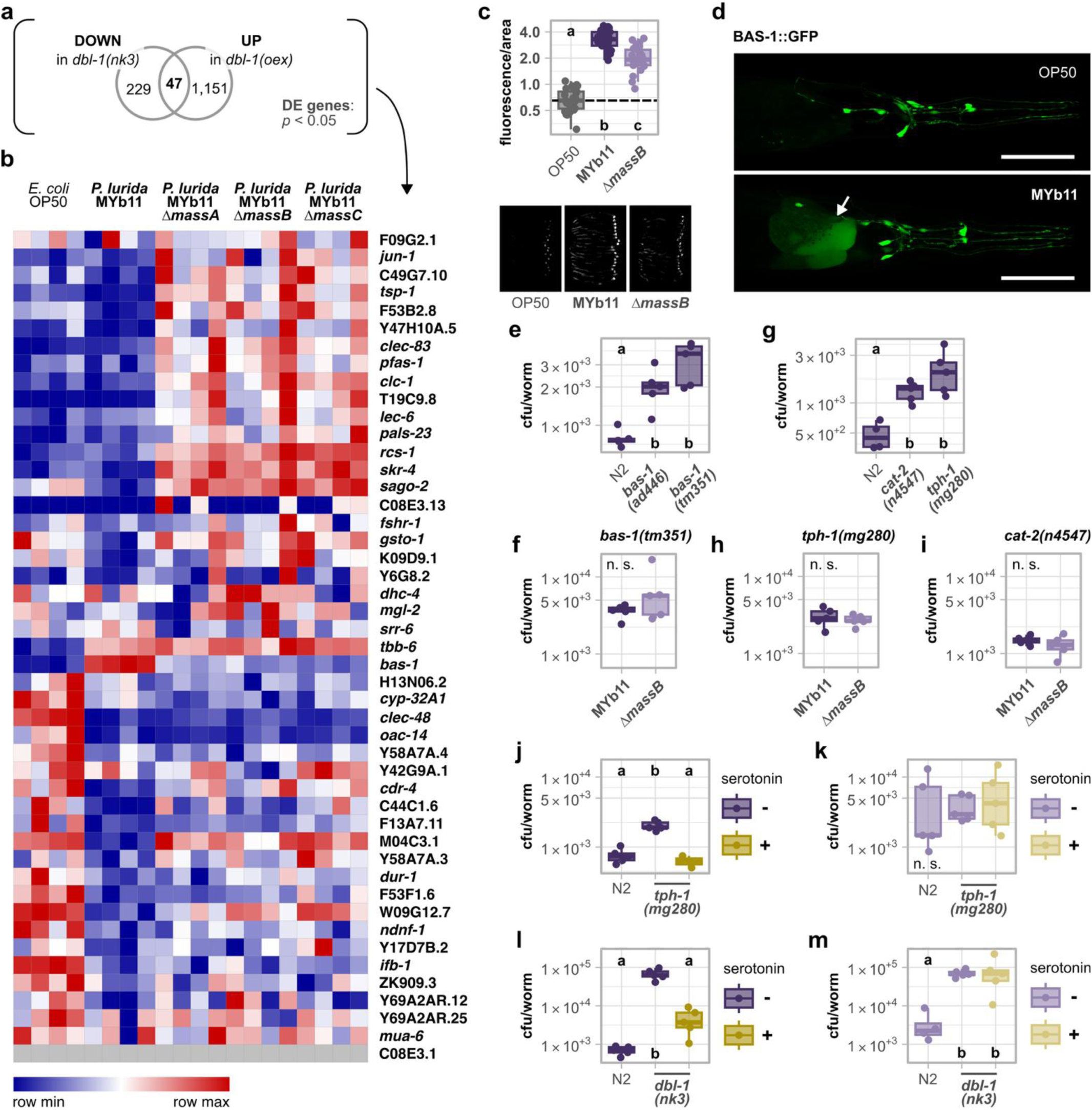
Serotonin acts downstream of DBL-1/TGF*-*β signaling to regulate *P. lurida* MYb11 colonization. (a) Venn diagram showing the number of significantly differentially expressed genes in the *dbl-1(nk)* mutant or the *dbl-1* overexpression compared to wildtype N2. Data taken from^29^. (b) Heatmap displaying the expression of genes positively regulated by TGF-β signaling, representing the overlap in Venn diagram. Worms were exposed to *E. coli* OP50, MYb11, or any of the massetolide-deficient MYb11 mutants, *ΔmassA*, *ΔmassB*, and *ΔmassC*. Shown are transcripts per million (TPM). Color intensity represents relative expression within each gene, with the color scale normalized to the minimum (blue) and maximum (red) TPM value for each row independently with n = 4. (c) Quantification of BAS-1::GFP fluorescence in 1 d adult transgenic worms exposed to OP50, MYb11, or MYb11 *ΔmassB*. Fluorescence was normalized by the worm’s body size (area). Each dot represents one worm with n = 25, and the dashed line represents the median of the mean gray value for OP50-exposed worms. Statistical significance (*p* ≤ 0.05) among groups of differently exposed worms was assessed using Kruskal–Wallis rank sum test followed by Dunn’s *post hoc* test with Holm correction. Different letters indicate significant differences, whereas shared letters indicate no significant difference. Representative images of groups of 20 individuals, arranged with their heads pointing to the right, are shown below. (d) Microscopic images of a transgenic strain carrying BAS-1::GFP (green) exposed to OP50 or MYb11. Shown are the head region including the first intestinal cells (indicated by the arrow) of 1 d adults. Scale bars represent 50 µm. (e-m) Bacterial load of MYb11 and MYb11 *ΔmassB*, in wildtype N2 or mutant worms. Plates were supplemented with (j-m) serotonin while exposure to (j, l) MYb11 or (k, m) MYb11 *ΔmassB*. Statistical significance (*p* ≤ 0.05) among treatment groups was assessed either using (e, g, j-m) the Kruskal–Wallis rank sum test followed by Dunn’s *post hoc* test with Holm correction or using (f, h, i) the Wilcoxon rank sum test with n = 3-5. Raw data and corresponding *p*-values are provided in Extended Data Table 1 and Supplementary Table 1, respectively.

To test if serotonin and/or dopamine act downstream of TGF-β signaling, we added exogenous serotonin and dopamine to the TGF-β *dbl-1(nk3)* mutant and assessed colonization by MYb11 wildtype and the massetolide-deficient *ΔmassB* mutant. Again, the high colonization phenotype of the *dbl-1* mutant was rescued by exogenous serotonin only for colonization by MYb11 wildtype (Figure 5l; Supplemental Figure 5d, e), not by the Δ*massB* mutant (Figure 5m). Yet, exogenous dopamine failed to rescue the colonization phenotype of the *dbl-1* mutant (Supplemental Figure 5f, g), indicating that either dopamine is not required downstream of TGF-β signaling for colonization control or that exogenous dopamine supplementation is not effective for restoring this phenotype. In contrast, the serotonin experiments consistently linked serotonergic signaling to TGF-β-dependent control of bacterial colonization in the presence of massetolide.

Taken together, these results imply that serotonin acts downstream of TGF-β signaling in the regulation of bacterial colonization, but only in the presence of massetolide. Exogenous serotonin cannot restore the high colonization phenotype in TGF-β *dbl-1* and serotonin biosynthesis *tph-1* mutants in the absence of the biosurfactant.

### Massetolide affects gut peristalsis via serotonin signaling and facilitates gut clearance through a low-surface-tension milieu

In *C. elegans*, the neurotransmitter serotonin is a key modulator of bacterial food-related behaviors and learned avoidance of pathogenic bacteria^31–35^. To better understand how serotonin functions in regulating colonization by MYb11, we first assessed the involvement of *C. elegans* behavior, in particular choice and feeding behavior, in the response to MYb11-derived massetolide. The cyclic lipopeptide biosurfactant serrawettin, which is produced by pathogenic *Serratia marcescens*, triggers a strong avoidance response, in which *C. elegans* initially moves onto the bacterial lawn, but later leaves it again^36^. However, *C. elegans* does not exhibit avoidance behavior when exposed to its symbiont MYb11. Thus, we assessed binary choice behavior. While *C. elegans* preferred its laboratory food bacterium *E. coli* OP50 over MYb11, it exhibited no choice preference between massetolide-producing wildtype MYb11 and non-producing mutants (Figure 6a; Supplemental Figure 6a). Next, we assessed pumping rate of worms on massetolide-producing MYb11 wildtype and massetolide-deficient MYb11 *ΔmassB* mutants. However, we did not observe any significant differences between the two bacterial strains (Figure 6b; Supplemental Figure 6b). Together, these findings indicate that MYb11-derived massetolide neither affects *C. elegans* choice, nor feeding behavior.

**Figure 6.**
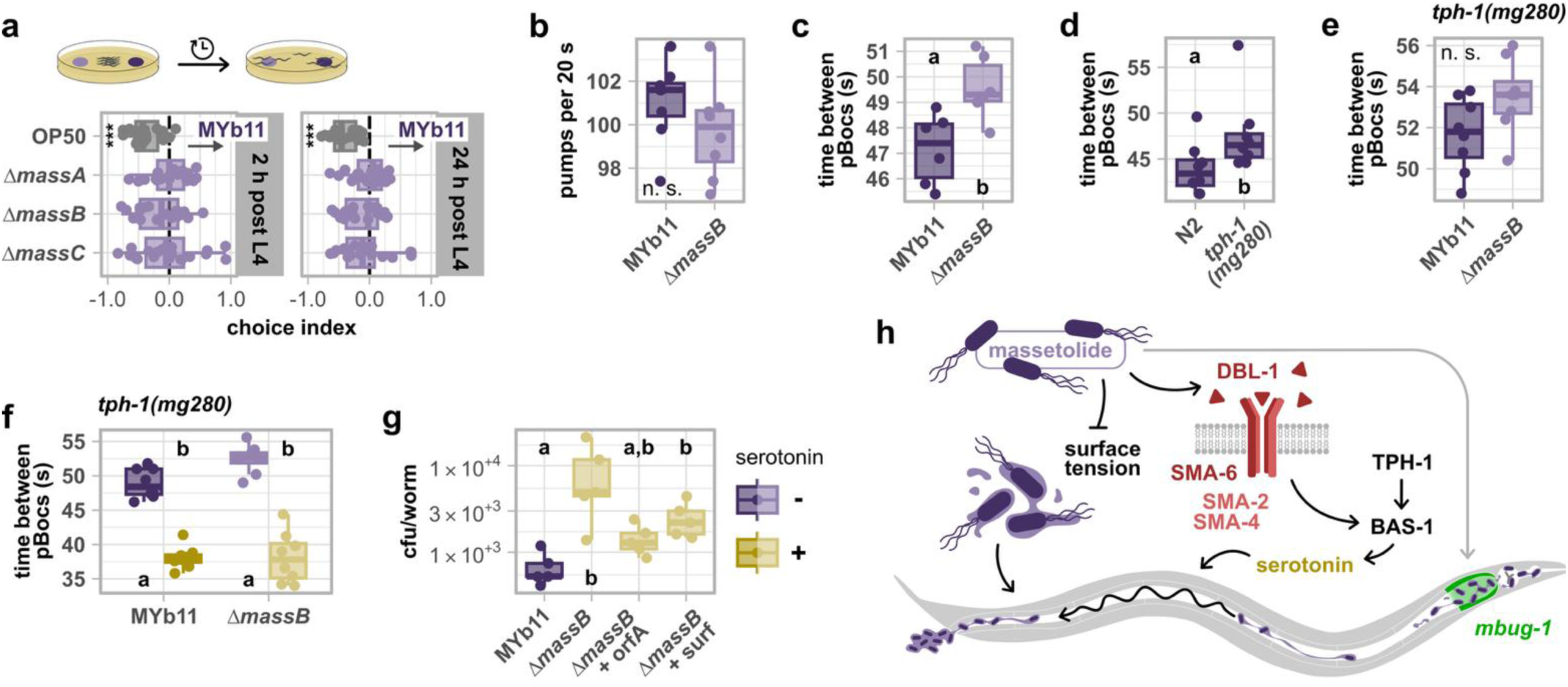
Massetolide affects gut peristalsis via serotonin signaling and enables gut clearance through a low-surface-tension milieu. (a) Choice behavior of wildtype N2 between *E. coli* OP50, massetolide-deficient MYb11 mutants, *ΔmassA*, *ΔmassB*, and *ΔmassC*, against MYb11. Choice index = (worms residing on MYb11 – worms residing on test bacterium) / worms residing on both bacterial spots. A negative choice index indicates a choice towards the test bacterium (y-axis), a positive choice index indicates a choice towards MYb11. Wilcoxon signed rank test per group against 0, FDR-corrected with n = 20, \*\*\**p* ≤ 0.001. Experimental set-up as schematic on top. (b) Ingestion rate of wildtype N2 on MYb11 and MYb11 *ΔmassB* measured as grinder movements (pumps) per 20 s in 1 d adults. (c-f) Defecation rate of (c, d) wildtype N2 and (d-f) *tph-1(mg280)* mutant on (c, e, f) both MYb11 and MYb11 *ΔmassB*, or (d) only MYb11 measured as time between two posterior body contractions (pBoc) in 1 d adults. (b-e) Statistical significance (*p* ≤ 0.05) among groups of differently exposed worms was assessed using the Wilcoxon rank sum test with (c) n =6 or (b, d, e) n = 8. (g) Bacterial load of MYb11 or MYb11 *ΔmassB* mixed with biosurfactants orfamide A (orfA) or surfactin (surf) in wildtype N2. MYb11 *ΔmassB* plates were supplemented with serotonin. (f, g) Statistical significance (*p* ≤ 0.05) among groups of differently exposed worms was assessed using the Kruskal–Wallis rank sum test followed by Dunn’s *post hoc* test with Holm correction with (f) n = 8 and (g) n = 5. (h) Model of MYb11 clearance through massetolide-induced neuroimmune signaling and reduction of surface tension. Raw data and corresponding *p*-values are provided in Extended Data Table 1 and Supplementary Table 1, respectively.

In mammals, serotonin is synthesized by enteric neurons and enteroendocrine cells within the gastrointestinal tract. Beyond its diverse physiological functions, serotonin plays a crucial role in modulating gut motility and contraction frequency^37^. Therefore, we asked if MYb11-derived massetolide influences the *C. elegans* defecation program. By measuring the time between posterior body contractions (pBocs), we found that the defecation frequency in *C. elegans* associated with massetolide-producing wildtype MYb11 was increased in comparison to worms exposed to non-producing MYb11 *ΔmassB* mutants (Figure 6c; Supplemental Figure 6c). These results indicate that massetolide increases intestinal peristalsis, which in turn decreases bacterial colonization. Indeed, in the serotonin biosynthesis mutant *tph-1(mg280)*, which showed a high colonization phenotype, defecation frequency was decreased on wildtype MYb11 in comparison to wildtype worms (Figure 6d). Importantly, we did not observe an effect of massetolide on gut peristalsis in the *tph-1(mg280)* mutant (Figure 6e, f), indicating that serotonin signaling mediates the effect of massetolide on peristalsis. Additionally, exogenous serotonin clearly increased defecation frequency in *tph-1(mg280)* mutants on both wildtype MYb11 and massetolide-deficient MYb11 *ΔmassB* mutants, so regardless of the MYb11 strain the host is associated with (Figure 6f). However, this increase in defecation frequency only led to a decrease in colonization when MYb11 produced massetolide (Figure 5j, l). As massetolide production reduces the surface tension in a MYb11 population (Figure 3a), the detergent-like milieu might facilitate bacterial clearance through gut peristalsis.

To test this hypothesis, we mimicked massetolide-mediated functions (i) by activating intestinal peristalsis using exogenous serotonin and (ii) by adding the cyclic lipopeptide biosurfactants orfamide A or surfactin – derived from *P. protegens* and *B. subtilis* – to the massetolide-deficient MYb11 *ΔmassB* mutant and measured bacterial load. Indeed, both biosurfactants partially reduced the bacterial load of the MYb11 *ΔmassB* (Figure 6g). Interestingly, orfamide A and surfactin also induced *mbug-1* expression similar to massetolide (Supplemental Figure 6d).

Taken together, massetolide induces host gut peristalsis through serotonin signaling and simultaneously creates the biophysical conditions favoring MYb11 expulsion (Figure 6h).

## Discussion

Our findings demonstrate that host control of symbiont abundance depends not only on host responses but also on microbial traits that determine susceptibility to those responses (Figure 6h). We identify the microbial molecule as well as the host signaling processes that determine colonization control in the *C. elegans*-MYb11 symbiosis. Moreover, we find that the same molecule renders MYb11 susceptible to that control. Massetolide increases serotonin-driven peristalsis, yet serotonin-dependent clearance is effective only against massetolide-producing bacteria. This suggests that host peristalsis selectively removes bacteria that possess particular physical properties, functioning as a mechanism for symbiont control rather than unspecific quantitative microbiota depletion. Consistent with this idea, gut transit time is increasingly recognized as an important determinant of microbiota composition in mammals^38–40^.

Our results identify the biosurfactant massetolide as a key determinant of this susceptibility. Beyond its established role in bacterial surface behavior and swarming motility, massetolide simultaneously acts as a host-directed signal that limits colonization. The importance of massetolide for host association is further supported by experimental evolution of *P. lurida* MYb11 toward host adaptation^41^. MYb11 is not directly transmitted vertically among host generations, but rather follows a bi-phasic life cycle where they alternate between a host associated and a free-living phase in the environment. This previous work demonstrated that swarming ability is advantageous in the free-living environmental phase and that, accordingly, host adaptation is accompanied by reduced swarming motility and downregulation of the massetolide biosynthetic gene cluster^41,42^. Similar effects of the cyclic lipopeptide biosurfactants surfactin and orfamide A suggest that the ability of bacterial biosurfactants to modulate host responses and intestinal colonization may represent a more general phenomenon. Together, these findings suggest that modulation of biosurfactant production may represent an adaptive strategy for tuning host colonization.

On the host side, we show that massetolide affects colonization through a host non-canonical DBL-1/TGF-β signaling pathway. How massetolide activates TGF-β signaling remains unclear. Given that biosurfactants can alter membrane and extracellular matrix (ECM) properties and that ECM remodeling can activate TGF-β pathways in *C. elegans*^43,44^, one possibility is that massetolide triggers signaling through physical perturbation of host surface structures. Future work will be required to determine how hosts sense biosurfactants and convert these signals into physiological responses.

The need for controlling symbiont abundance may be particularly important for symbionts such as *P. lurida* MYb11 that exhibit context-dependent effects on host fitness. While MYb11 can protect against infection, excessive proliferation is detrimental to the host^9–11,20^. Mechanisms that restrict bacterial abundance without eliminating the symbiont may therefore help maintain a beneficial association while limiting potential costs. More broadly, our findings show how a single microbial metabolite can simultaneously promote host-mediated clearance by peristalsis and render specific bacteria biophysically susceptible to that clearance, thereby coupling host and microbial physiology to regulate symbiont abundance.

## Materials and Methods

### Strains, maintenance, and preparations

Wildtype *C. elegans* N2 and all used *C. elegans* mutants and transgenic strains, as well as the bacterial strains, were received from sources indicated in Supplementary Table 2. Worm strains were maintained according to standard procedures^45^. For each experiment, worms were synchronized by bleaching gravid hermaphrodites with alkaline hypochlorite solution and incubating the eggs in M9 overnight on a shaker.

Spore solutions of pathogenic *Bacillus thuringiensis* strains MYBt18247 (Bt247) and MYBt18679 (Bt679) and stock solutions of non-pathogenic Bt407 were prepared following a previously established protocol^46^, and stored at -20 °C. Single aliquots were freshly thawed for each experiment.

*Pseudomonas lurida* MYb11 belongs to the natural microbiota of *C. elegans*^18^ and is stored in glycerol stocks at -80 °C. Before each experiment, bacterial isolates were streaked from glycerol stocks onto TSB (tryptic soy broth) agar plates, grown for 2 d at 25 °C, and consequently for an overnight in TSB at 28 °C in a shaking incubator. Nematode growth medium (NGM) plates were inoculated with the bacterial culture and incubated at 25 °C for an overnight before use.

### Transposon insertion mutant library generation

Transposon insertion mutant libraries with 11,047 mutants of *P. lurida* MYb11 were generated using the EZ-Tn5TM <R6Kγori/KAN-2>Tnp Transposome Kit (LGC Biosearch Technologies, Hoddesdon, United Kingdom). For this purpose, MYb11 were inoculated in LB medium and incubated overnight at 30 °C and 120 rpm in a shaking incubator. Overnight cultures were pelleted at 6 mL per reaction, the supernatant removed, and cells washed three times with 1 mL of 300 mM sucrose. Cells were resuspended and aliquoted in 100 µL of 300 mM sucrose. Fresh aliquots were placed on ice and 0.5 µL of transposome added per aliquot. 500 ng of the plasmid pME6032 served as positive control and an empty aliquot as negative control. Each aliquot was electroporated in a pre-cooled 2 mm cuvette using a Bio-Rad Gene-Pulser (Feldkirchen, Germany) at 25 µF, 200 Ω and 2.5 kV. 900 µl of prewarmed SOC medium were immediately added to the cells, which were incubated for 1 h at 30 °C and 120 rpm. After incubation, 100 µL of undiluted cells were plated onto LB agar plates supplemented with 30 µg/mL kanamycin (LB-Kan; GERBU Biotechnik, Heidelberg, Germany). The plates were incubated for 24 h, after which individual colonies were picked into 96 well plates filled with 150 µl LB-Kan. After 24 h of incubation the mutant libraries were stored by overlaying 50 µL of LB-Kan and DMSO (Carl Roth, Karlsruhe, Germany) at a final concentration of 8%. The mutant libraries were frozen and stored at -80 °C.

To verify transposon insertion into the *P. lurida* MYb11 genome and check for the location, mutants were picked at random for insertion site screening using a rescue-cloning approach. Genomic DNA was extracted from MYb11 mutants using the Wizard Genomic DNA Extraction Kit (Promega, Madison, WI, USA) according to manufacturer’s instructions for Gram-negative bacteria and eluted in 50 µL ddH2O. The extracted gDNA was restricted using 5 µL of PstI-HF und 5 µL cutsmart buffer (NEB, Ipswich, MA, USA) in a 50 µL reaction for 15 min at 37 °C. The reaction was then purified using the NucleoSpin Gel and PCR Clean-up kit (Macherey-Nagel), according to manufacturers’ instructions. The restricted DNA was self-ligated by adding 2U of T4 DNA Polymerase and 2 µL Ligase Buffer (Thermo Scientific), which were incubated for 1 h at room temperature. The reaction was heat inactivated for 10 min at 70 °C and 3 µL heat-shock transformed into competent *E. coli* DH5α at 42 °C for 90 s. The cells were subsequently incubated on ice for 5 min. 800 µL of SOC-medium was added to the cells, which were incubated for 1 h at 37 °C and shaking. Transformed *E. coli* were plated onto LB-agar with kanamycin (30 µg/mL). After overnight incubation at 37 °C, colonies were picked and self-ligated plasmids extracted using the PrestoTM Mini Plasmid kit (Geneaid, New Taipei City, Taiwan) according to manufacturers’ instructions. Plasmid DNA was amplified by PCR using the transposon specific primers Kan-2-FP1-For and R6Kan-2-RP-1-Rev supplied by the transposome kit (LGC Biosearch Technologies). The PCR reaction included 2 µL template, 5 µL 5x GoTaq Green Buffer, 2.5 µL MgCl2 (25 mM), 1.5 µL dNTPs, 1 µL of each primer Kan-2-FP1-For and R6Kan-2-RP-1-Rev at 10 mM, 0.25 µL GoTaq Polymerase (Promega) and nuclease-free water added to a volume of 25 µL. Primer targets were amplified in a GeneTouch Thermocycler (Bioer, Hangzhou, China) for 1 cycle 3 min and 95 °C, 25 cycles of 30 s at 94 °C, 30 s at 57 °C and 2.5 min at 72 °C, and a final elongation for 5 min at 72 °C. PCR products were purified using the NucleoSpin Gel and PCR Clean-up kit (Macherey-Nagel) and sent for Sanger sequencing at MicroSynth (Göttingen, Germany). Sequences provided the flanking regions to transposon inserts, which were mapped onto the *P. lurida* MYb11 genome to identify the insertion region using the software Unipro UGENE^47^.

### Worm imaging and quantification

For imaging of *in vivo* gene expression, transgenic 1 d adults (1 d post L4) and 4 d adults (4 d post L4) were anesthetized with 10 mM tetramisole, placed onto slides containing a fresh 2% agarose patch, and imaged with a Leica stereomicroscope M205 FA (Wetzlar, Germany). Magnification and exposure time for the fluorophore signal were kept the same in each experiment to ensure comparability; contrast and brightness were adjusted for representative images (grouped worms).

Gene expression of reporter strains was quantified using ImageJ2 version 2.16.0/1.54p^48^. Worms were individually imaged and the integrated density (IntDen) of each worm was measured. To correct for potential worm size differences IntDen values were normalized by the total area of each respective individual.

For microscopy, worms were mounted on slides with a dried 2% agarose patch and immobilized with 10 mM tetramisole. All pictures were taken using the Zeiss confocal microscope LSM 700 (Jena, Germany).

### *P. lurida* MYb11 transposon mutant library screen

Wells of 96-well plates were prepared with 150 µL agarose NGM (agarose replacing agar agar). *P. lurida* MYb11 transposon mutants were grown in round bottom 96-well plates, each well containing 100 µL TSB with 30 µg/mL kanamycin, shaking at 28 °C overnight. Each agarose NGM well was inoculated with 100 µL of overnight culture and left at 25 °C another overnight. Then, small L1 populations of synchronized *mbug-1*p::GFP were added to each well and kept at 20 °C. After one day, worms were manually screened under a fluorescence stereomicroscope for failed induction of *mbug-1*. Empty wells without any bacteria and inevitably no worm development were noted. Positive *P. lurida* MYb11 transposon mutants were thoroughly evaluated a second time to exclude false positives.

The transposon insertion site of positive candidates was determined by rescue cloning using the manufacturer’s protocol (EZ-Tn5^TM^ <R6Kγori/KAN-2>Tnp Transposome^TM^ Kit, epicentre, Cat. No. TSM08KR; Madison, WI, USA).

### Generation of *P. lurida* MYb11 knock-out mutants and GFP-tagged MYb11 *ΔmassB*

A two-step allelic exchange protocol was applied in order to generate *P. lurida* MYb11 knock-out mutants^49^. ∼700 bp PCR amplicons flanking each desired gene deletion were cloned into the pUIsacB suicide plasmid and transformed into competent *E. coli* cells. The constructed plasmid was transferred into MYb11 recipients by tri-parental conjugation using a helper *E. coli* strain containing plasmid pRK2013^50^. Successful transconjugants were first selected by tetracycline and nitrofurantoin and then counter-selected using sucrose medium. Presence of the deletion mutations was confirmed by Sanger sequencing.

Tagging *P. lurida* MYb11 mutant *ΔmassB* with sfGFP followed established protocols using an *E. coli* donor strain carrying pTn7xKS-sfGFP with the kill switch^51^. The identity of obtained fluorescent isolate by PCR was confirmed with *ΔmassB* specific primers SHP1733 and SHP1734, the integration of sfGFP with insert-specific primers WP11 and WP256 (Supplementary Table 3). A list of all genetically modified *P. lurida* MYb11 is to be found in Supplementary Table 2.

### Swarming and drop collapse assay, *in vitro* antimicrobial activity, and growth dynamics

Overnight cultures of *P. lurida* MYb11 and *P. lurida* MYb11 *Δmass* mutants were adjusted with 1x PBS (phosphate-buffered saline) to OD_600_ of 1. 2 µL of each bacterial suspension was dropped onto soft agar plates (TSB with 0.6% agar agar) and incubated at 20 °C. Bacterial growth on the plates was imaged after 4 d.

To demonstrate the presence or absence of the water surface tension in bacterial liquid cultures, bacterial overnight cultures were dropped onto parafilm and immediately photographed.

To assess antimicrobial activity of *P. lurida* MYb11 *Δmass* mutants, an overnight culture of *B. thuringiensis* Bt679 in TSB was adjusted to an OD_600_ of 10, diluted 1:100 into soft agar (1% agar in TSB) cooled to 55 °C, and poured into 9 cm diameter plates. After several hours, 2 µL of each overnight culture of the focal bacteria were spotted onto the Bt–soft agar plates and incubated for at least 24 h at 20 °C.

To measure bacterial growth dynamics, TSB overnight cultures were adjusted to an OD_600_ of 0.5 using 1x PBS and inoculated into liquid NGM (standard NGM without agar) at a 1:20 dilution. Bacterial growth was monitored by recording OD_600_ every 30 min for 24 h using a plate-reader (Epoch 2, Agilent BioTek, Winooski, VT, USA) with continuous shaking at room temperature.

### Infection, survival and lifespan experiments

Generally, synchronized L1 larvae were grown on NGM plates with overnight lawns of the focal bacteria including *E. coli* OP50 as control at 20 °C.

Infection with *Pseudomonas aeruginosa* PA14 followed established slow-killing protocols^52^.

For survival experiments, peptone-free medium (PFM, nematode growth medium without peptone) plates for Bt infection were prepared the day before adding L4 larvae: bacteria of an overnight culture were harvested by centrifugation, resuspended in 1x PBS, pH7, adjusted to OD_600_ of 10, serially diluted with Bt spores, and eventually used for plate inoculation. As L4s, worms were rinsed off the plates and washed with M9 and pipetted in populations of approximately 30 worms on each Bt infection plate. After 24 h incubation at 20 °C survival of worms was scored. Worms were considered to be alive when they moved upon gentle prodding with a worm pick. Replicates with less than 15 worms at the time of scoring were excluded.

For lifespan experiments, 30 synchronized L4 larvae were transferred onto NGM plates seeded with overnight cultures of the focal bacteria. Worm survival was assessed daily, and live adults were regularly transferred to fresh plates with the same bacterial treatment until the end of the egg-laying period.

### Bacterial load experiments

To measure bacterial load, 1 d adult worms were rinsed off the plates and repeatedly washed with 0.025% Triton-X in M9 (M9+T), then surface sterilized with 2% sodium hypochlorite solution in M9+T (NaClO; Roth, 9062.3), washed again with 0.025% Triton-X in PBS (PBS+T), pH7. A distinct number of worms (approx. 10-25) was ground with Zirconia beads in a Bead Ruptor 96 (Omni International; Kennesaw, GA, USA) at 30 Hz for 3 min. Bacterial solutions were diluted, plated onto TSB plates, and colony forming units (cfu) were scored after 1 d at 20 °C. The cfu count of the supernatant which was taken before worm beating was subtracted from the final cfu count.

### RNA-seq analyses

The published RNA-seq data describing TGF-β signaling targets was available under the accession number GSE186653^29^. To identify *dbl-1*-acivated target genes, the kallisto output files were fed into the DESeq2 workflow^53^ and genes with *p*_adj_ < 0.05 were considered as differentially expressed (Extended Data Table 1).

To assess gene expression in response to massetolide, wildtype N2 were synchronized and grown on NGM seeded with overnight cultures of either *E. coli* OP50, *P. lurida* MYb11, or *P. lurida* MYb11 *Δmass* mutants at 20 °C. The populations were gravity washed with M9 and transferred to new plates each day of adulthood in order to separate the adult worms from their offspring. The experiment was conducted with four independent biological replicates. For RNA isolation, 1 d adults (24 h post L4) and 4 d adults (96 h post L4) were processed by snap-freezing washed worm pellets in TRI Reagent (Zymo Research, Cat. No. R2050-1-200) and five freeze-thaw cycles in liquid nitrogen and at 45 °C. RNA was extracted using the Direct-zol RNA MiniPrep kit (Zymo Research, Cat. No. R2052).

Stranded mRNA libraries were generated with the DNBSEQ (MGI, Shenzhen, Guangdong, China) workflow by BGI (Yantian, Shenzhen, China): Total RNA was poly(A)-enriched using oligo(dT) magnetic beads, fragmented, and reverse-transcribed with random hexamer primers to synthesize first-strand cDNA; second-strand synthesis incorporated dUTP in place of dTTP. The double-stranded cDNA underwent end repair, 3′ A-tailing, and adaptor ligation. To retain strand specificity, the dUTP-containing strand was removed with uracil-DNA glycosylase, and the remaining strand was amplified by limited-cycle PCR. Amplified products were heat-denatured, circularized with a splint oligonucleotide and DNA ligase, and converted into DNA nanoballs via rolling-circle amplification, which were subsequently sequenced on the DNBSEQ platform with paired-end 150 bp length.

After quality control, raw reads were filtered using SOAPnuke^54,55^ removing adaptors, reads with N content greater than 1%, and low quality reads (if more than 40% of their bases with quality score ≤ 20). The resulting clean reads were mapped to the *C. elegans* reference genome WBcel235 using HISAT2 (v2.0.4^56^) to account for splicing. Subsequently, Bowtie2 (v2.2.5^57^) was used to align the same reads to reference transcriptome to obtain gene-level alignment results for expression quantification. RSEM^58^ then estimated transcript-level expression, resulting in TPM, FPKM, and expected count values. Differential gene expression was determined according to a DESeq2^53^ workflow including pre-filtering that only kept genes with >= 10 reads in at least 4 samples (n = 4). The threshold for significant differential gene expression was set to *p*_adj_ < 0.001 and a log_2_ fold-change > 1. The heatmap showing expression patterns in *dbl-1*-activated target genes was created using TPM with the online tool Morpheus (https://software.broadinstitute.org/morpheus).

### Binary choice, ingestion and defecation experiments

Plates for the choice experiment were prepared by spotting 30 µL of two test bacteria overnight cultures opposite of each other onto 6 cm diameter plates, and incubating them at 25 °C overnight. 30-50 naïve L4 larvae were pipetted centrally between both bacterial spots. The number of worms residing on each bacterium was determined after 2 and 24 h. The choice index was calculated as [(number of worms on bacterium A − number of worms on bacterium B) / total number of worms]. To assess ingestion and defecation, worms were grown on the focal bacteria to 1 d adults. The number of grinder movements was counted over 20 s, and the interval between two successive posterior body contractions (pBoc) was measured, respectively.

### Neurotransmitter and biosurfactant supplementation

Serotonin hydrochloride (8367.3, Carl Roth, Karlsruhe, Germany) was freshly prepared in 0.1 M HCl at a concentration of 1 M, and dopamine hydrochloride (H8502, Sigma-Aldrich/Merck, Darmstadt, Germany) in M9 buffer at 50 mM. NGM plates were supplemented with serotonin to a final concentration of 5 mM no later than one day before bacterial inoculation. Control plates were supplemented with an equivalent volume of 0.1 M HCl. Inoculated NGM plates (approximately 10 mL agar per plate) were treated with 400 µL of dopamine solution or an equal volume of M9 as a control immediately prior to worm transfer following^59^. To ensure continuous dopamine exposure, L4-stage worms were transferred to freshly prepared dopamine-containing plates at the L4 stage.

Massetolide F (kindly provided by Ashootosh Tripathi at Natural Products Discovery Core, University of Michigan Life Sciences Institute; Ann Arbor, MI, USA), orfamide A (AA Blocks, Cat. No. AA01E5UC; San Diego, CA, USA), and surfactin (TargetMol, T13041; Boston, MA, USA) were dissolved in DMSO to a working stock of 20 µg/µL. To supplement overnight bacterial cultures with biosurfactants, cultures were centrifuged, and a defined volume of the supernatant was removed and replaced with an equal volume of the appropriate biosurfactant solution or DMSO as a control to reach a massetolide F concentration of 0.2 µg/µL or 2 µg/µL in the inoculum, or an orfamide A or surfactin concentration of 0.4 µg/µL in the inoculum.

### Statistical analyses

All remaining statistical analyses were carried out with RStudio, R v4.2.1, graphs created with its package ggplot2 v3.3.6^60^, and edited with Inkscape v1.1.2. Statistics per dataset are listed in Supplementary Table 1 and 4.

## Data availability

Strains and plasmids generated can be made available on request. RNA-seq data has been deposited to ENA’s ArrayExpress with the accession E-MTAB-17622. Source data are provided with this paper.

## Declaration of generative AI and AI-assisted technologies in the manuscript preparation process

During the preparation of this work, the authors used ChatGPT (OpenAI) for language editing and as a conversational tool for critically examining the clarity, coherence, and persuasiveness of arguments and their presentation. The authors reviewed and edited the output as needed and take full responsibility for the content of the published article.

## Acknowledgments

We are grateful to Nirmal Chaudhary and Ashootosh Tripathi from the Natural Products Discovery Core at Michigan University for providing purified massetolide F, the Nick Burton lab at Van Andel Institute in Grand Rapids, MI, USA, for their collaboration, Sabrina Butze for technical support, and the Schulenburg group particularly Hinrich Schulenburg for their valuable feedback on the project. We also thank Herman Vargas Gebauer for his technical support during the generation and verification of the transposon mutant library, as well as Hendrik Bethge and Finn Reimers for their support with transposon mutant colony picking. We further thank the Caenorhabditis Genetics Center (University of Minnesota, Minneapolis, Minnesota, USA), funded by the NIH Office of Research Infrastructure Programs (P40OD010440) for *C. elegans* strains. Additionally, some strains were provided by NBRP, which is funded by the Japanese government.

## Author contributions

BP and KD conceived the study, designed the experiments, interpreted the data, and wrote the manuscript. BP performed the experiments and analyzed the data. CEH conducted and analyzed some TGF-β experiments, AHH together with RAS generated and verified the MYb11 transposon mutant library. BP, KD, RAS, and PR secured the funding. All authors reviewed and approved the final manuscript.

## Supplemental Figures

**Supplemental Figure 1.**
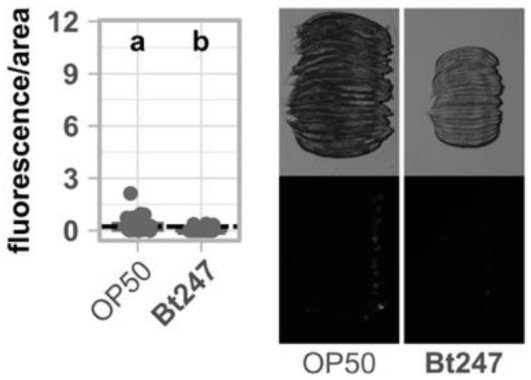
*B. thuringiensis* Bt247 does not induce *mbug-1* expression. Quantification of *mbug-1*p::GFP fluorescence in 1 d adult transgenic worms exposed to *E. coli* OP50 and pathogenic *B. thuringiensis* Bt247. Fluorescence was normalized by the worm’s body size (area). Each dot represents one worm with n = 24–32, and the dashed line represents the median of the mean gray value for OP50-exposed worms. Statistical significance (p ≤ 0.05) among groups of differently exposed worms was assessed using the Wilcoxon rank sum test. Different letters indicate significant differences, whereas shared letters indicate no significant difference. Representative images of groups of 20 individuals, arranged with their heads pointing to the right, are shown below in bright field (top) and fluorescence (bottom). Raw data and corresponding *p*-values are provided in Extended Data Table 1 and Supplementary Table 4, respectively.

**Supplemental Figure 2.**
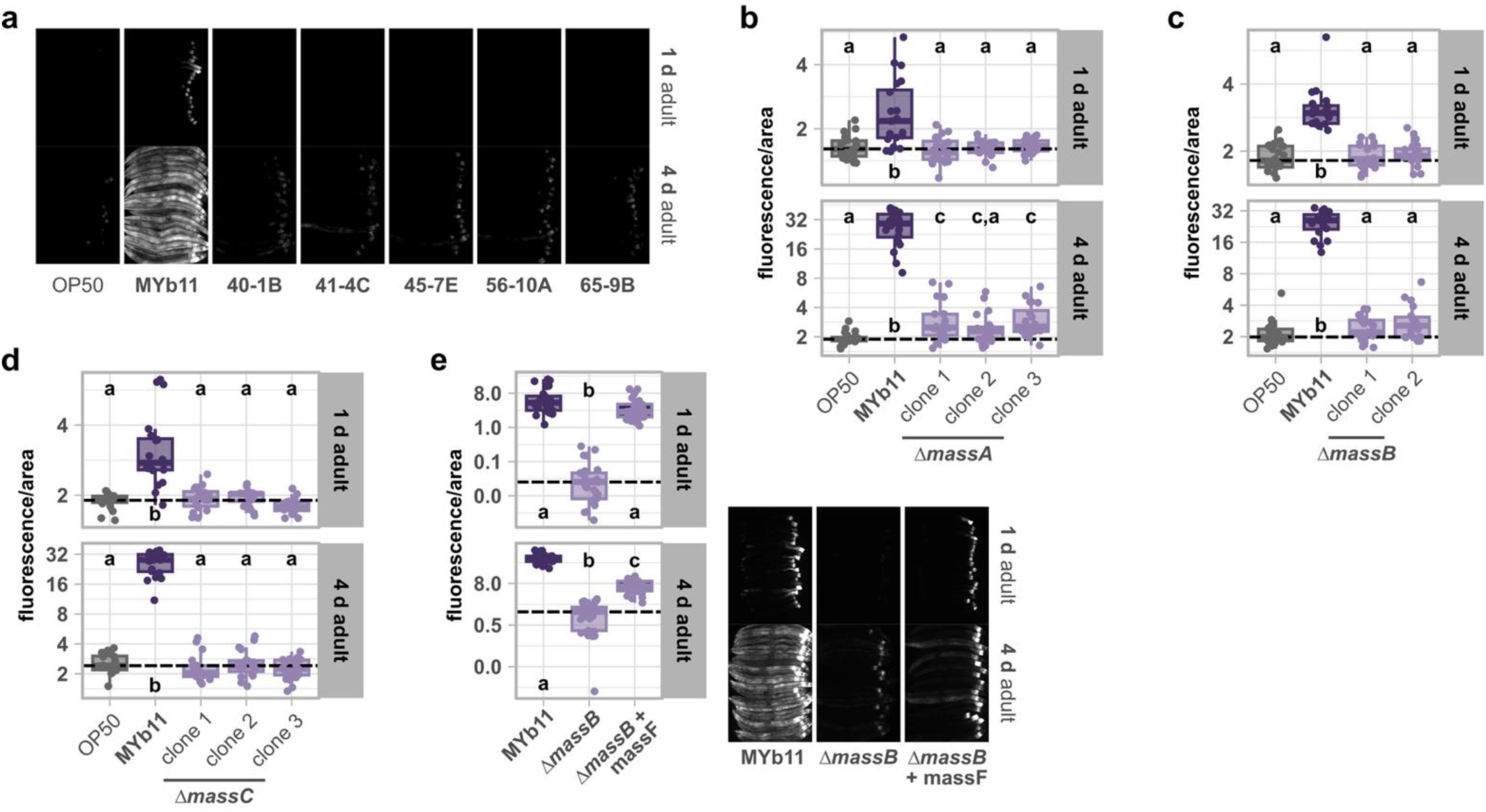
*P. lurida* MYb11 transposon mutants and massetolide-deficient mutants fail to induce *mbug-1* expression. (a) Expression of *mbug-1*p::GFP in 1 d and 4 d adults exposed to *E. coli* OP50, *P. lurida* MYb11, or the five MYb11 transposon mutants derived from the screen. Representative fluorescence images of groups of 20 individuals, arranged with their heads pointing to the right, are shown. (b-e) Quantification of *mbug-1*p::GFP fluorescence in 1 d and 4 d adult transgenic worms exposed to OP50, MYb11, or MYb11 *Δmass* single mutants, or (e) MYb11, MYb11 *ΔmassB*, or MYb11 *ΔmassB* supplemented with 0.2 µg/µL purified massetolide F in the inoculum. Fluorescence was normalized by the worm’s body size (area). Each dot represents one worm with n = 25, and the dashed line represents the median of the mean gray value for (b-d) OP50-exposed or (e) MYb11-exposed worms. Statistical significance (*p* ≤ 0.05) among groups of differently exposed worms was assessed using the Kruskal–Wallis rank sum test followed by Dunn’s *post hoc* test with Holm correction. Different letters indicate significant differences, whereas shared letters indicate no significant difference. Raw data and corresponding *p*-values are provided in Extended Data Table 1 and Supplementary Table 4, respectively.

**Supplemental Figure 3.**
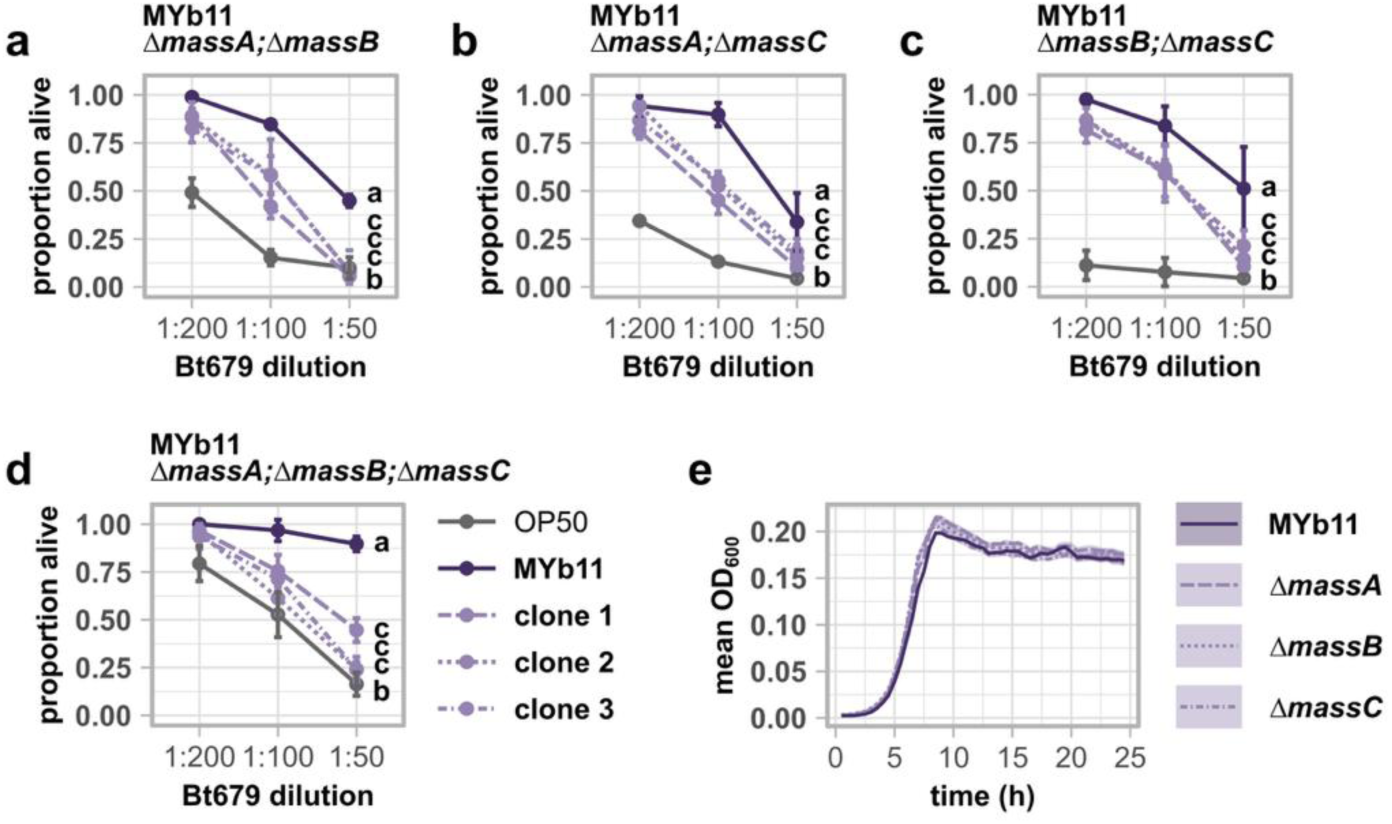
*P. lurida* MYb11-mediated protection against *B. thuringiensis* Bt679 infection is partially reduced in worms exposed to *P. lurida* MYb11 *Δmass* double/triple mutants. (a-d) Survival of wildtype N2 infected with serial dilutions of *B. thuringiensis* Bt679 after 24 hours post infection (hpi). Worms were fed with either *E. coli* OP50, *P. lurida* MYb11, or clones of massetolide (a-c) double and (d) triple mutants before and during infection. Each dot represents the mean ± standard deviation of three worm populations, n = 3. The same letters indicate no significant differences between the dose-response curves according to a generalized linear model followed by Tukey-adjusted *post hoc* multiple comparisons. (e) Growth curves of *P. lurida* MYb11 and the MYb11 *Δmass* mutants. The lines represent the mean of n = 16 technical replicates, the shading represents the standard error. Raw data and corresponding *p*-values are provided in Extended Data Table 1 and Supplementary Table 4, respectively.

**Supplemental Figure 4.**
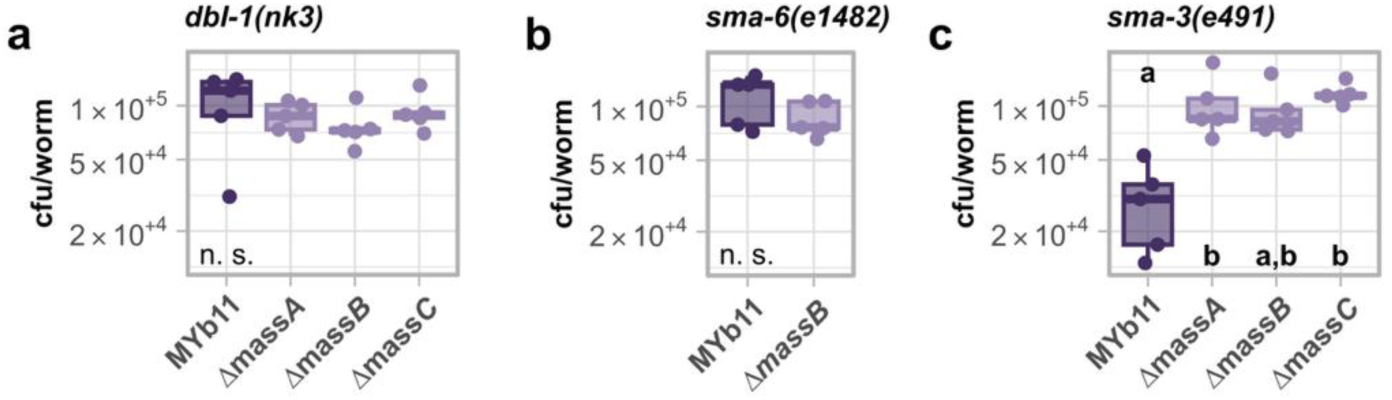
Massetolide affects *P. lurida* MYb11 colonization through DBL-1/TGF-β signaling. Bacterial load of *P. lurida* MYb11 and MYb11 *Δmass* mutants in TGF-β signaling mutants. Statistical significance (*p* ≤ 0.05) among treatment groups was assessed either using (a, c) the Kruskal–Wallis rank sum test followed by Dunn’s *post hoc* test with Holm correction or using (b) the Wilcoxon rank sum test with n = 5. Raw data and corresponding *p*-values are provided in Extended Data Table 1 and Supplementary Table 4, respectively.

**Supplemental Figure 5.**
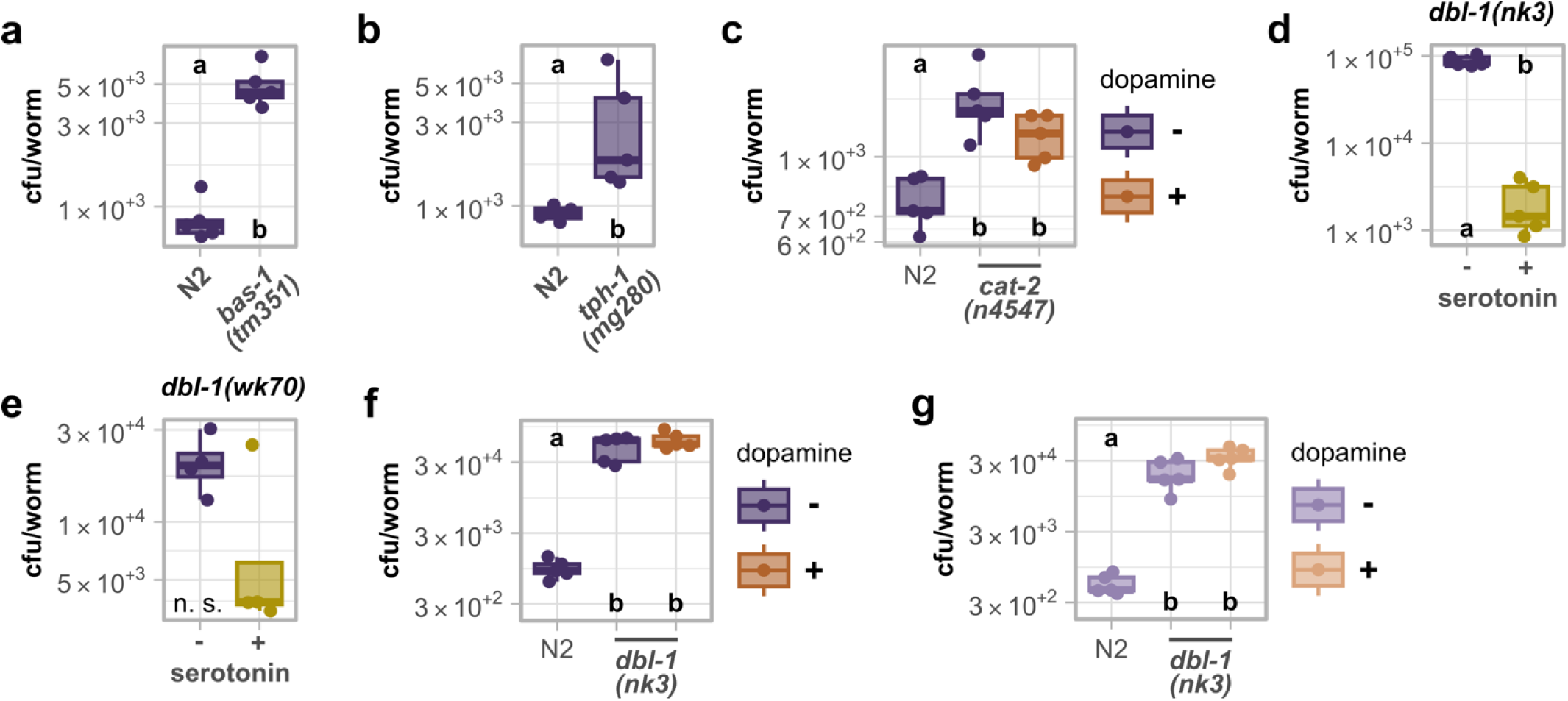
Serotonin acts downstream of DBL-1/TGF-β signaling to regulate *P. lurida* MYb11 colonization. Bacterial load of (a-f) *P. lurida* MYb11 or (g) MYb11 *ΔmassB* in wildtype N2 or mutant worms. Plates were supplemented with (d, e) serotonin or (c, f, g) dopamine while exposure to (c-f) MYb11 or (g) MYb11 *ΔmassB*. Statistical significance (*p* ≤ 0.05) among treatment groups was assessed either using (a, b, d, e) the Wilcoxon rank sum test or using (c, f, g) the Kruskal–Wallis rank sum test followed by Dunn’s *post hoc* test with Holm correction, n = 4-5. Raw data and corresponding *p*-values are provided in Extended Data Table 1 and Supplementary Table 4, respectively.

**Supplemental Figure 6.**
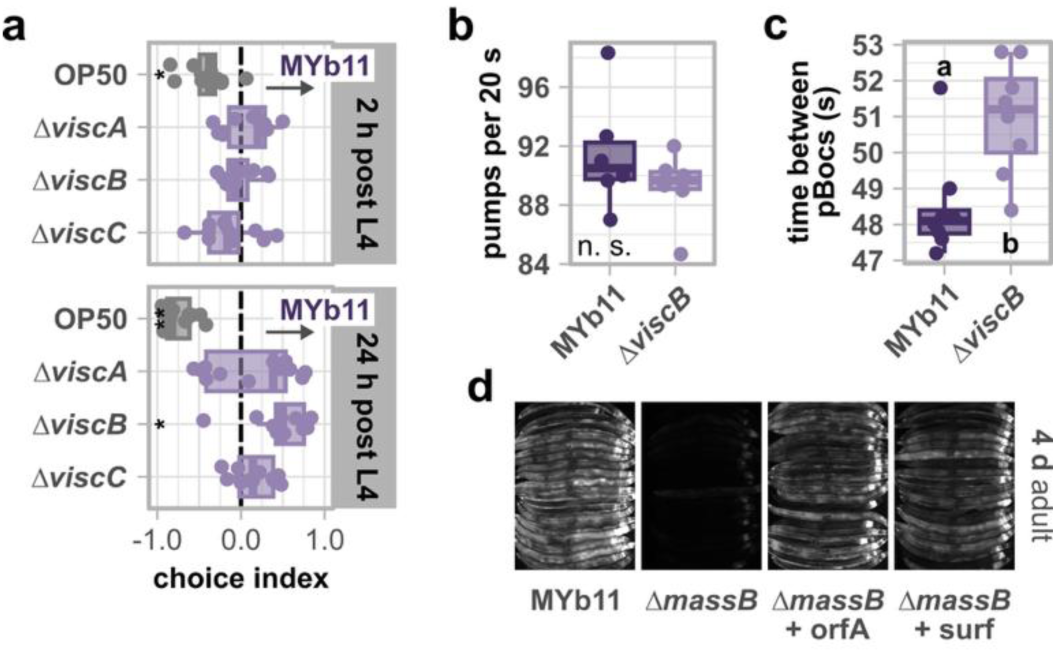
Massetolide affects gut peristalsis and biosurfactants orfamide A and surfactin induce *mbug-1* expression. (a) Choice behavior of wildtype N2 between *E. coli* OP50, massetolide-deficient MYb11 mutants, *ΔmassA*, *ΔmassB*, and *ΔmassC*, against MYb11. Choice index = (worms residing on MYb11 – worms residing on test bacterium) / worms residing on both bacterial spots. A negative choice index indicates a choice towards the test bacterium (y-axis), a positive choice index indicates a choice towards MYb11. Wilcoxon signed rank test per group against 0, FDR-corrected with n = 11-13, \**p* ≤ 0.05, \*\**p* ≤ 0.01. (b) Ingestion rate of wildtype N2 on MYb11 and MYb11 *ΔmassB* measured as grinder movements (pumps) per 20 s in 1 d adults. (c) Defecation rate of wildtype N2 on MYb11 and MYb11 *ΔmassB* measured as time between two posterior body contractions (pBoc) in 1 d adults. (b, c) Statistical significance (*p* ≤ 0.05) among groups of differently exposed worms was assessed using the Wilcoxon rank sum test with (b) n = 6 and (c) n = 8. (d) Expression of *mbug-1*p::GFP in 4 d adults exposed to MYb11, MYb11 *ΔmassB*, or MYb11 *ΔmassB* supplemented with biosurfactants orfamide A (orfA) or surfactin (surf). Fluorescence images of groups of 20 individuals, arranged with their heads pointing to the right, are shown. Raw data and corresponding *p*-values are provided in Extended Data Table 1 and Supplementary Table 4, respectively.

